# msaGUI: Multispectral Analysis Graphical User Interface for Ratiometric Analysis and Background Correction

**DOI:** 10.64898/2026.06.30.735666

**Authors:** Grayson R. Hoy, Caitlin M. Davis

## Abstract

Chemical imaging is a powerful branch of modern microscopy encumbered by a lack of flexible, high-throughput analysis tools. Bespoke analytical pipelines typically perform ratiometric analysis on two layers in a multispectral image to describe the relative composition of molecules in a sample. This strategy has been implemented across fields, spanning histopathology, cell biology, environmental science, and materials science. The commercialization of chemical imaging microscopes has facilitated the collection of large multispectral datasets, necessitating accessible ways to process them. This paper describes Multispectral Analysis Graphical User Interface (msaGUI), a desktop graphical user interface to analyze individual and batch datasets of multispectral images. Data is loaded as CSV, TSV, or TIFFs and processed through a user-defined sequence of modular image operations that can be flexibly combined, e.g. to reduce spectral crosstalk or background noise. After analysis, data is visualized as exportable images, histograms, and statistics. To yield publication-quality figures, outputted images are fully customizable. Written in Python with open-source libraries, the msaGUI program is packaged into an executable for Windows and Mac for a fully no-code application. Other operating systems are supported via the Python source code. In summary, msaGUI provides a rapid and user-friendly solution for analyzing and visualizing multispectral data.

## Introduction

As technology and manufacturing have advanced, commercial infrared and Raman microscopes have become more accessible and affordable. Microscopes are available from numerous companies, including Bruker, DRS Daylight Solutions, Oxford Instruments, Horiba, and JASCO, among others. These companies offer a variety of chemical imaging microscopes designed for diverse applications, such as compact designs, high-resolution imaging, and other specialized capabilities. Additionally, companies are rapidly implementing state-of-the-art approaches that would traditionally be limited to a specialist. For example, Leica’s STELLARIS Coherent Raman Scattering microscope can operate in stimulated Raman scattering, coherent anti-stokes Raman scattering, and second harmonic generation modalities. Photothermal Spectroscopy Corp. has commercialized sub-micron Optical Photothermal Infrared (O-PTIR) technology. These commercially available microscopes are user-friendly, with intuitive interfaces and simplified operation facilitating the rapid collection of high volumes of data.

By detecting vibrations, chemical imaging techniques provide spectroscopic fingerprints capable of identifying molecules in a sample without labels. In environmental and materials science, chemical imaging has revealed the origin of microplastics based on its composition and tested the efficiency of semiconductors.^[1,2]^ Chemical imaging is also powerful in biomedical research; it can distinguish pathological tissues and monitor cellular dynamics.^[3,4]^ However, these techniques generate dense data sets that often require postprocessing to resolve the features of interest.^[5–7]^ Each single frequency image reveals important spatial features unique to molecules with a vibrational mode at that frequency. Spectral demixing and normalization of images is achieved through comparative analysis, which is frequently accomplished by dividing different spectral images into ratios.^[8–10]^

Despite the potential of chemical imaging methods, the large volume of data generated can create a bottleneck in postprocessing. This necessitates the development of high-throughput analytical tools capable of efficiently parsing and processing chemical images. Specialized microscopy software exists for such needs, but open-source options remain limited. Programs that are compatible across instrument brands are increasingly important as chemical imaging becomes more commercialized. Although image analysis exists for the open-source software Fiji, it lacks the batch visualization necessary for large sets of experiments.^[11]^ Additionally, image correction for noise and extraneous signal requires several steps. The current landscape reveals a gap: the lack of open-source software capable of flexible batch analysis and visualization of images, which includes tools such as correcting for background noise and spectral crosstalk. To close this gap, we developed an open-source, multiplatform software solution that allows for dynamic batch analysis and visualization. This software is designed to be both intuitive and easily visualizable, allowing for complex analysis regardless of instrument brand. Further, it provides a framework for batch processing of more complex multi- and hyperspectral analyses in the future.

## Results and Discussion

### Graphical User Interface and Output

msaGUI is implemented in Python 3 using the tkinter library; however, Python or associated libraries are not required for use. The program is downloadable via executables specific to each operating system. At publication, it is available for Windows and Mac. It was tested on Windows 11 and Mac Sonoma and Tahoe. For all operating systems, the raw Python code is available on Github. The main window presented at startup allows the user to add images and specify a sequence of image operations to be completed in batch (Figure 1). After analysis, generated images and histograms are available in the designated export folder with a customizable file type (e.g. JPEG, PNG, EPS). Histograms are included to supplement the images’ spatial information with quantitative descriptions of intensity distribution. Statistics for each data file can be exported into a single spreadsheet for simple batch analysis.

**Figure 1.**
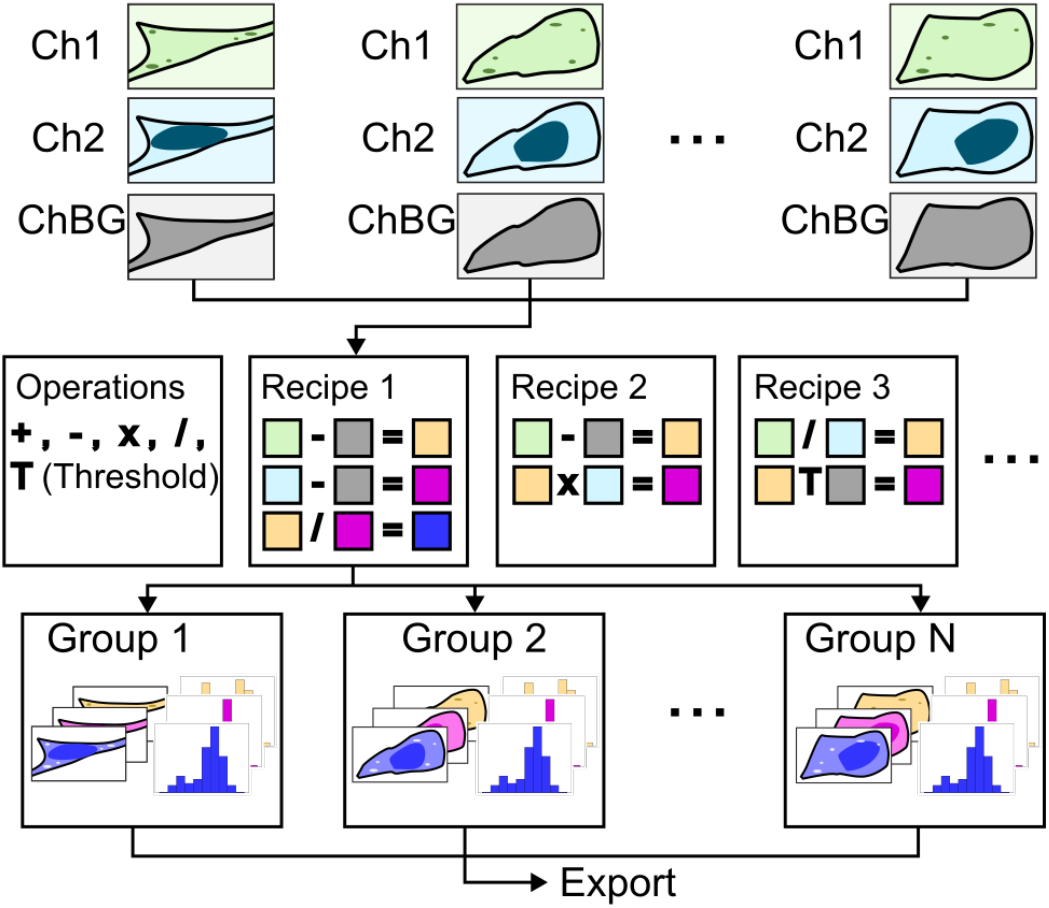
msaGUI analysis workflow. User inputs images for an arbitrary number of image groups and sets up analysis pipeline. The analysis pipeline is comprised of a modular set of operations with user defined recipes. The batch analysis is performed, yielding processed images, histograms, and associated statistics of each group. If desired, these outputs are exported to a filesystem outside of the program.

### Code Availability

The msaGUI is available in the GitHub repository located at https://github.com/cdavislab/multispectral-analysis. Users are encouraged to report issues, suggest features, and contribute code improvements. A tutorial and guide (GUIDE.pdf) is included in the repository, with installation instructions, usage guidelines, and a tutorial. msaGUI analyses including Image Correction, Thresholding, Ratio Imaging, and Statistics are described below. msaGUI uses the following open-source libraries: NumPy^[12]^, Matplotlib^[13]^, Pillow^[14]^, SciPy^[15]^, H5Py^[16]^, and Pytest^[17]^.

### Image Correction

Image correction removes unwanted confounding signals from the images. Factors such as sample background, optical errors, and detector response can be corrected by acquiring an image without the sample and subtracting it from the sample image. Applying such an approach, as opposed to a uniform correction, provides flexibility with how each pixel is corrected. Further, some samples require pixel-specific spectral demixing to isolate the signal of interest. If such flexibility is not necessary, simple constant value subtraction is also available.

More complicated corrections are often necessary. To create images that accurately reflect the absorption of the molecule of interest, these correction factors must be determined and used to demix the multifrequency images. In homogenous samples, when two or more vibrational transitions in a molecule’s spectrum occur at or very close to the same frequency, it is impossible to distinguish them using a single frequency image. The characteristic contribution of each peak to the frequency of interest can be determined by fitting the molecule’s full spectrum to a combination of the appropriate functions for the sample (e.g. Gaussian, Lorentzian, Voigt) (Figure 2). In heterogeneous samples, the measured total intensity in each pixel is a linear combination of the absorptions of the individual molecule(s) within the voxel. The intensity of the overlap depends on the concentration of the overlapping molecule(s).^[18]^ In such cases, the characteristic percentage of the overlapping bands at the frequency of interest in relation to a band that is unique to each molecule can be determined in a full spectrum of each molecule. An image must be collected at one of the unique frequencies so that the image at the frequency of interest can be corrected.

**Figure 2.**
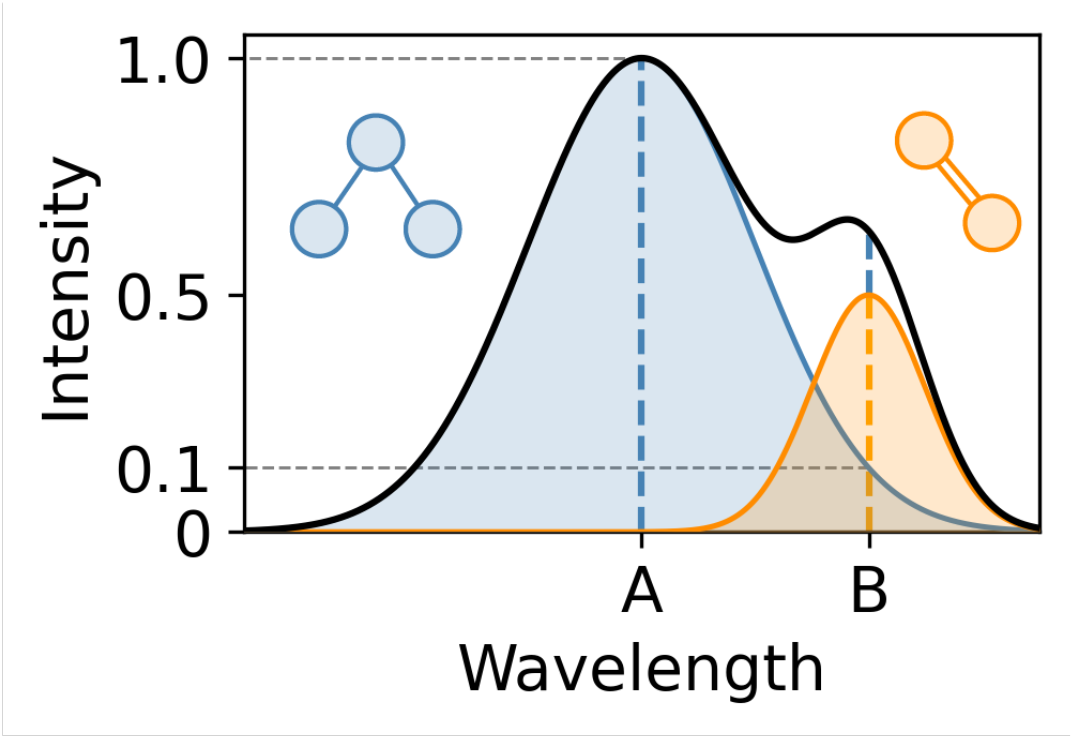
Signal from two unique molecules (blue and orange) sum together in a single spectrum (black). Signal from one molecule (blue) is completely resolvable at wavelength A. Signal from another molecule (orange) is not resolvable with a single wavelength. Fitting the spectrum (black) to a combination of Gaussians deconvolves the individual blue and orange contributions. Unmixing shows that 10% of blue signal at A is at B. Since only the blue molecule is present at A, subtracting 10% of the total signal at A from B reveals the true signal of the orange molecule at B.

The msaGUI analysis setup allows users to construct a series of arithmetic image operations, providing the ability to correct images with a known correction factor and a correction mask image collected at the associated correction frequency. The program allows for the subtraction of overlap in each image:

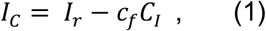

where *I*_*C*_ is the corrected image, *I*_*r*_ is the raw image, *c*_*f*_ is the correction factor, and *C*_*I*_ is the correction image. Correction is per pixel, and each pixel is treated independently. This style of image correction can be used with background images by setting the correction factor to 1 or for pixel-specific spectral demixing using the pre-determined correction factor related to the correction intensity image to reveal the desired signal.

### Thresholding

To remove background noise that would otherwise affect the output visualizations and statistics, thresholding is optionally available. However, it should be noted that thresholding is subjective and could remove real low intensity image information.

In the msaGUI, the threshold is input as (1) a proportion (*p*) of the maximum value (*I*_*max*_) in the image, (2) a set value (*V*), or (3) another image (*I*_*2*_). Mathematically,

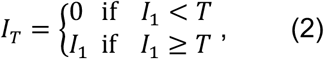

where *I*_*T*_ is the thresholded image, *I*_1_ is the input image, and *T* is the threshold (1) *pI*_*max*_, (2) *V*, or (3) *I*_2_. Thresholds are evaluated per pixel. For example, in case (1) an image with a max value of 100 and threshold of 0.05 would result in all pixel values less than or equal to 5 being set to not a number (NaN) in the ratio image and excluded from statistics calculations. Any pixel that is NaN will be visualized with a distinct color.

### Ratio Imaging

A ratio of images at two frequencies offers several advantages over a single frequency image. The ratio can correct for variations in signal intensity across the sample arising from factors such as sample thickness and surface variation.^[19]^ Ratios effectively normalize the concentration providing improved chemical contrast. For the same reason, ratioing can reduce background noise. Comparing the relative intensity, it may be possible to distinguish between a signal of interest and background.^[20]^ Finally, in some cases the ratio of specific vibrational modes are correlated with specific chemical properties. For example, degree of crystallinity in carbon materials^[21,22]^ or distinguishing healthy and diseased tissue.^[23,24]^

As an option in the user-defined pipeline, images in msaGUI can be ratioed together pixel-by-pixel with:

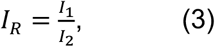

where *I*_*R*_ is the ratio image, *I*_1_ is the image in the numerator, and *I*_2_ is the image in the denominator. In cases where a pixel value in *I*_2_ = 0, the corresponding value in *I*_*R*_ is set to not a number (NaN) to avoid an undefined quotient. In some contexts, it may be desired to add a small constant prior to thresholding to avoid NaN values.^[25,26]^

### Statistics

The mean 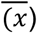, median 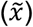, and maximum (*x*_*max*_) intensities for all raw and processed images are calculated in addition to standard deviation (*σ*), standard error (*SE*), and total number of pixels (*n*). Formal definitions can be found in the Supplementary Information.

### GUI Operation

When the user opens the GUI, they are presented with several panels and inputs (Figure 3). Initially empty, the file list (Figure 3A) displays files that are added by the user or generated during analysis. To load files, the “Add Files” button (Figure 3C) prompts a file dialog box, where the user can select multiple files at the same time. The program accepts comma-delimited (CSV), tab-delimited (TSV), and TIFF files. For CSV and TSV files, each value represents a pixel value from the image (Figure S1). Once added, files are represented in the file list as their file name. Optionally, file names can be displayed with their parent directory or full file path. After selecting a file in the file list, it is immediately viewable as images (Figure 3B) or histograms. The View submenu on the menu bar at the top of the window has a variety of options to change how and which files are represented in the file list. Data can be viewed together in groups and shown as images or histograms. It can also be sorted based on name, group, time imported, or keyword. Further, images with specific keywords can be hidden to organize data. Files can be shown as just the file name, with their parent folder, or as their full file path. Removing files can be easily done by selecting a file and clicking the “Delete Files” button (Figure 3C). Once files are selected, all analysis is performed using the “Analyze” button (Figure 3C).

**Figure 3.**
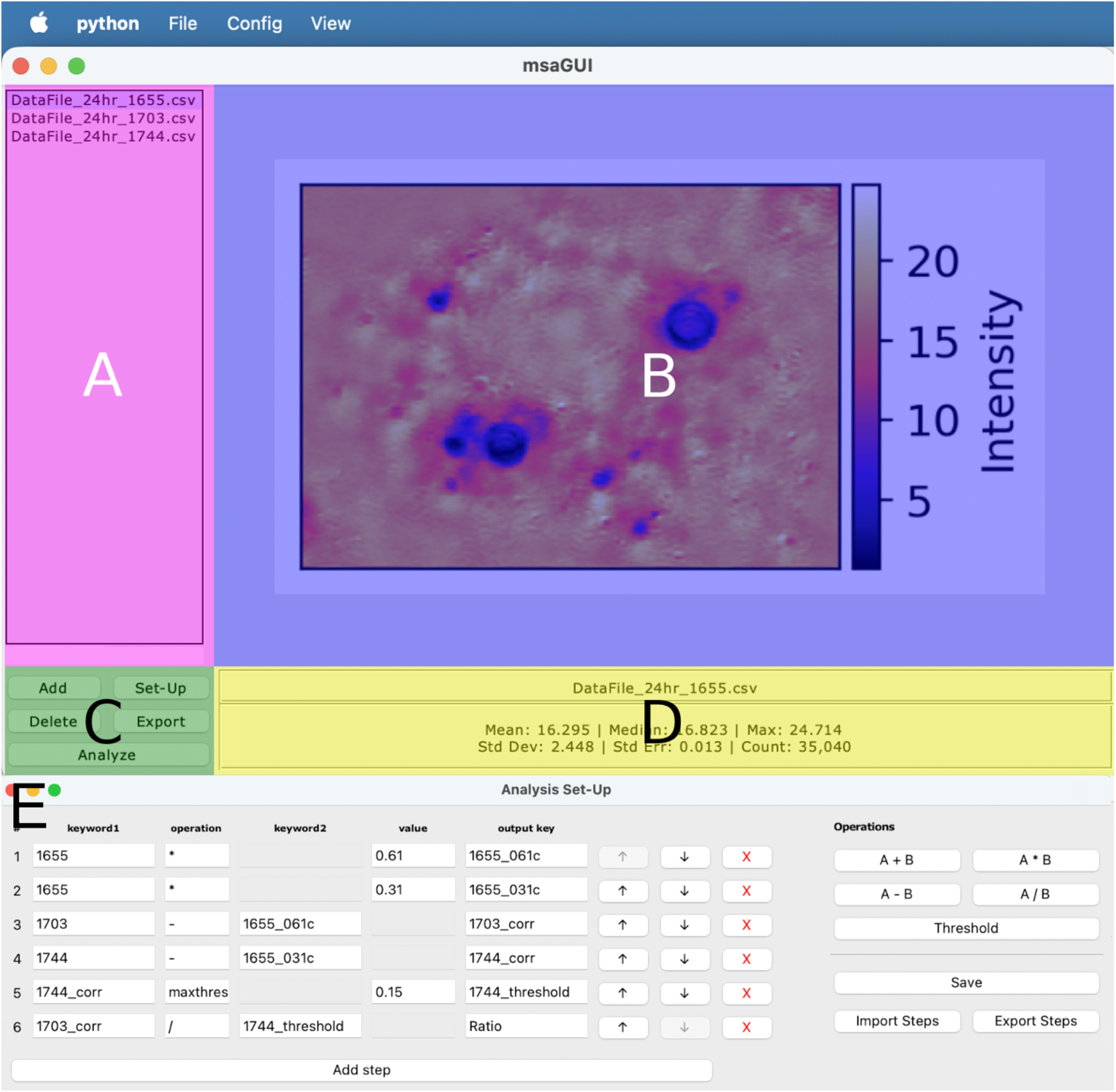
msaGUI interface false colored and labeled by feature: (A) file list (magenta), (B) data visualizer (blue), (C) file buttons (green), (D) file descriptors (yellow) and (E) analysis set-up menu (grey). Clicking the (C) Set-Up button opens (E) the analysis set-up menu. In the analysis set-up menu users can input steps on the left by typing in a keyword, operation, output key, and either a second keyword or value. Alternatively, users can input steps with the operation buttons on the right side. Steps can be rearranged or deleted with the up, down, and x buttons to the side of each step. A series of modular operations (i.e. an analysis pipeline) can be exported and later imported with the buttons on the right side. Save confirms the set-up for analysis.

To enable batch processing, msaGUI matches images together for ratioing and correction based on their file names. The program assumes that files in the same group are identically named except for a difference in keywords inputted to the GUI. These keywords describe the different channels used to collect the data (e.g., a frequency, “1660 cm^−1^”). Keywords are defined alongside the modular operations in the Set-Up panel (Figure 3E). Files of the same image (i.e. different layers) must have the same folder path. For example, the program will group files “C://Dir/Experiment1_1744” and “C://Dir/Experiment1_1703” together if 1703 and 1744 are input in either the keyword1 or keyword2 column. In the setup described in Figure 3E, first the 1703 and 1744 are background subtracted with a 1655 image. Then a 0.1 threshold is applied to the corrected 1744 image. Finally, the corrected 1703 image is divided by the corrected and thresholded 1744 image.

Once adjustable parameters are set (Figure 3E), clicking the “Analyze Files” button groups (Figure 3C), corrects, thresholds, and ratios the inputted files. Each ratioed set of data produces a corresponding image and histogram. All images in a single group can be visualized at once in a group view, toggleable via the View submenu. To resize the viewer, the border between the fie list and viewer can be moved to prioritize the viewer, and the entire window can be resized. The adjustable parameters can be imported and exported using the File subfolder on the menu bar. All items generated by the analysis are saved in one folder, which can be defined explicitly during export (Figure 3C). If an export folder is not specified, items will be automatically saved to the subfolder /msa_analysis where the code is run. For convenience, the files can be further separated into folders based on the original data’s parent folder name. An example output folder path is “/msa_analysis/Expt1/DataFile1”.

### Use Case: Tracking *De Novo* Lipogenesis in Huh-7

We present a case of msaGUI used to determine the rate of storage of *de novo* synthesized triglycerides and cholesterol esters into lipid droplets in Huh-7 cells (gift from Lars Plate, Vanderbilt University) via O-PTIR. The experiment has been previously described.^[4]^ Briefly, cells were grown and plated onto CaF_2_ (Crystran, Poole, U.K) coverslips in Dulbecco’s Modified Eagle Medium (DMEM, Corning, Corning, NY) with 10% fetal bovine serum and 1% penicillin-streptomycin (Thermo-Fischer, Waltham, MA). After 24 hours, the media was replaced with a glucose-free DMEM supplemented with 10% fetal bovine serum and 1% penicillin-streptomycin, 4.5 mg/mL ^13^C-glucose (D-glucose-^13^C_6_, CLM-1396-PK, Cambridge Isotope Laboratories), and 2% fatty acid-free bovine serum albumin (BSA, Sigma-Aldrich, St. Louis, MO). At 24, 48, and 72 hours post ^13^C-glucose feeding, Huh-7 cells were imaged with O-PTIR at three different frequencies: 1655 cm^−1^, 1703 cm^−1^, and 1744 cm^−1^. 1655 cm^−1^ arises from water background and the amide I band. 1703 cm^−1^ and 1744 cm^−1^ arise from the ^13^C=O and ^12^C=O triglyceride/cholesterol ester carbonyl stretches, respectively. Correction factors for spectral demixing were calculated by collecting full IR spectra outside of lipids of ^13^C-glucose and ^12^C-glucose fed Huh-7 cells. An example of calculating these correction coefficients is available (Figure S2). Briefly, the images were normalized to 1655 cm^−1^ at the water/amide I mode and the average percent of overlap of the water/amide I band at 1703 cm^−1^ or 1744 cm^−1^ was determined.

The average relative intensities between 1744 cm^−1^ and 1655 cm^−1^ in these low lipid areas yielded a correction coefficient of 0.61. The same procedure identified the 1703 cm^−1^ correction coefficient to be 0.31. The collected images, stored as grayscale CSV files, were inputted into msaGUI and processed according to Figure 3E. The program then automatically analyzed the 96 files corresponding to 32 cells in batch, saved and outputted images, histograms, and statistics (Figure 4A-E). Outputs were condensed for publication; see raw outputs in Figure S3-S4. Using the modular operations in msaGUI, the analysis in Castillo et. al was rapidly replicated (Figure 4F).^[4]^

**Figure 4.**
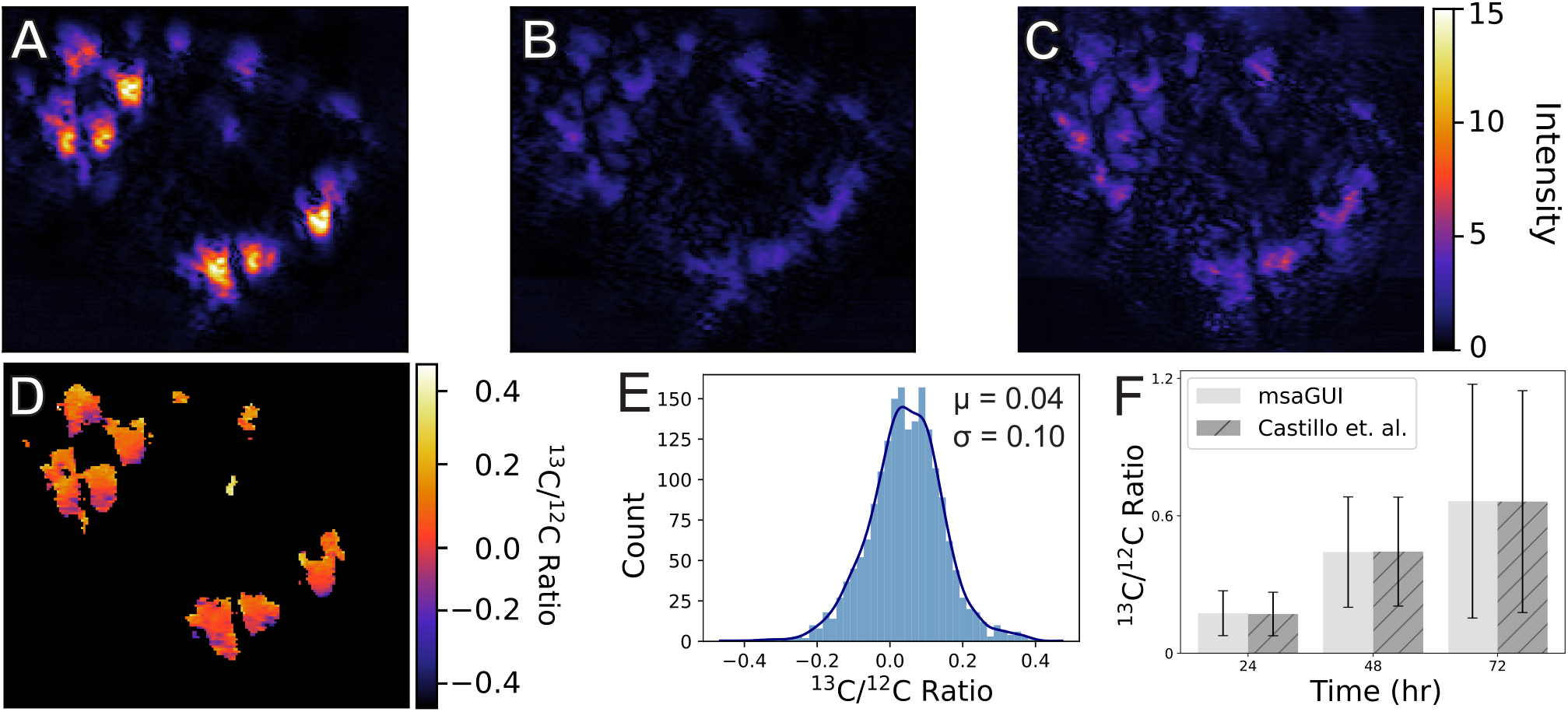
O-PTIR image of a Huh-7 cell 72 hours after feeding with ^13^C-glucose collected at (A) 1744 cm^−1^, (B) 1703 cm^−1^, and (C) 1655 cm^−1^. (D) Ratio image of 1703 cm^−1^ and 1744 cm^−1^, both corrected for overlapping signal at 1655 cm^−1^. (E) The ratios can also be displayed as a histogram. (F) Comparison of results between msaGUI and previously published results.^[4]^

## Conclusion

msaGUI is a robust, open-source software designed to expedite the analysis of multispectral microscopy data. We illustrated its efficacy in processing and visualizing complex imaging datasets with the case study of *de novo* lipogenesis in Huh-7 hepatocytes using O-PTIR. The software’s ability to adjust for spectral crosstalk and background noise, as well as its ratiometric imaging calculations, make it particularly useful for characterizing molecular composition and viewing snapshots of cellular processes. While we present an application in cell biology, the msaGUI will be broadly useful in vibrational imaging applications.

The software’s modular design enables it to handle a wide range of experimental setups, making it a useful tool for a variety of researchers. msaGUI simplifies the analytical workflow, allowing for rapid processing and intuitive data presentation and analysis. Moreover, as an open-source software requiring no coding experience but enabling bulk analysis and visualization of datasets, msaGUI fills a niche in the existing image analysis environment.

To summarize, msaGUI provides an effective, user-friendly platform to simplify high-throughput data analysis. We invite the scientific community to adopt and contribute to the continuous development of this tool, so that it can continue to meet the changing needs of researchers.

## Supporting Information

Code is available at https://github.com/cdavislab/multispectral-analysis. Included is example data used to produce the case study and a comprehensive tutorial for using the example and other data. Additional supporting information can be found online in the Supporting Information.6/30/26 6:24:00 PM

## Acknowledgements

This work was supported by a Beckman Young Investigator Award from the Arnold and Mabel Beckman Foundation (https://dx.doi.org/10.13039/100000997). G.R.H. was partially supported by a National Science Foundation Graduate Research Fellowship (DGE-2139841). The authors thank Dr. Michael Burke, Hannah Castillo, Anna Curtis, and Sydney Shuster for testing and user feedback. Microsoft Copilot was used for copyediting.

## References

[1] D. Schymanski, C. Goldbeck, H.-U. Humpf, P. Fürst, “Analysis of microplastics in water by micro-Raman spectroscopy: Release of plastic particles from different packaging into mineral water” Water Res. 2018, 129, 154–162.

[2] Y. G. Gogotsi, V. Domnich, S. N. Dub, A. Kailer, K. G. Nickel, “Cyclic nanoindentation and Raman microspectroscopy study of phase transformations in semiconductors” J. Mater. Res. 2000, 15, 871–879.

[3] S. Kumar, T. Verma, R. Mukherjee, F. Ariese, K. Somasundaram, S. Umapathy, “Raman and infra-red microspectroscopy: towards quantitative evaluation for clinical research by ratiometric analysis” Chem. Soc. Rev. 2016, 45, 1879–1900.

[4] H. B. Castillo, S. O. Shuster, L. H. Tarekegn, C. M. Davis, “Oleic acid differentially affects lipid droplet storage of de novo synthesized lipids in hepatocytes and adipocytes” Chem. Commun. 2024, 60, 3138–3141.

[5] S. O. Shuster, A. E. Curtis, C. M. Davis, “Optical Photothermal Infrared Imaging Using Metabolic Probes in Biological Systems” Anal. Chem. 2025, 97, 8202–8212.

[6] S. O. Shuster, M. J. Burke, C. M. Davis, “Spatiotemporal Heterogeneity of De Novo Lipogenesis in Fixed and Living Single Cells” J. Phys. Chem. B 2023, 127, 2918–2926.

[7] H. B. Castillo, C. M. Davis, “Optical photothermal infrared imaging of fatty acid esterification in the ER of living cells” Sci. Adv. in press.

[8] X. Huang, J. Song, B. C. Yung, X. Huang, Y. Xiong, X. Chen, “Ratiometric optical nanoprobes enable accurate molecular detection and imaging” Chem. Soc. Rev. 2018, 47, 2873–2920.

[9] L. Wu, C. Huang, B. P. Emery, A. C. Sedgwick, S. D. Bull, X.-P. He, H. Tian, J. Yoon, J. L. Sessler, T. D. James, “Förster resonance energy transfer (FRET)-based small-molecule sensors and imaging agents” Chem. Soc. Rev. 2020, 49, 5110–5139.

[10] S. Al Jedani, C. Lima, C. I. Smith, P. J. Gunning, R. J. Shaw, S. D. Barrett, A. Triantafyllou, J. M. Risk, R. Goodacre, P. Weightman, “An optical photothermal infrared investigation of lymph nodal metastases of oral squamous cell carcinoma” Sci. Rep. 2024, 14, 16050.

[11] J. Schindelin, I. Arganda-Carreras, E. Frise, V. Kaynig, M. Longair, T. Pietzsch, S. Preibisch, C. Rueden, S. Saalfeld, B. Schmid, J.-Y. Tinevez, D. J. White, V. Hartenstein, K. Eliceiri, P. Tomancak, A. Cardona, “Fiji: an open-source platform for biological-image analysis” Nat. Methods 2012, 9, 676–682.

[12] C. R. Harris, K. J. Millman, S. J. van der Walt, R. Gommers, P. Virtanen, D. Cournapeau, E. Wieser, J. Taylor, S. Berg, N. J. Smith, R. Kern, M. Picus, S. Hoyer, M. H. van Kerkwijk, M. Brett, A. Haldane, J. F. del Río, M. Wiebe, P. Peterson, P. Gérard-Marchant, K. Sheppard, T. Reddy, W. Weckesser, H. Abbasi, C. Gohlke, T. E. Oliphant, “Array programming with NumPy” Nature 2020, 585, 357–362.

[13] J. D. Hunter, “Matplotlib: A 2D Graphics Environment” Comput. Sci. Eng. 2007, 9, 90–95.

[14] “Pillow,” can be found under https://pillow.readthedocs.io/en/stable/index.html (accessed 28 March 2026), n.d.

[15] P. Virtanen, R. Gommers, T. E. Oliphant, M. Haberland, T. Reddy, D. Cournapeau, E. Burovski, P. Peterson, W. Weckesser,J. Bright, S. J. van der Walt, M. Brett, J. Wilson, K. J. Millman, N. Mayorov, A. R. J. Nelson, E. Jones, R. Kern, E. Larson, C. J. Carey, İ. Polat, Y. Feng, E. W. Moore, J. VanderPlas, D. Laxalde, J. Perktold, R. Cimrman, I. Henriksen, E. A. Quintero, C. R. Harris, A. M. Archibald, A. H. Ribeiro, F. Pedregosa, P. van Mulbregt, “SciPy 1.0: fundamental algorithms for scientific computing in Python” Nat. Methods 2020, 17, 261–272.

[16] “HDF5 for Python — h5py 3.16.0 documentation,” can be found under https://docs.h5py.org/en/stable/ (accessed 28 March 2026), n.d.

[17] H. Krekel, B. Oliveira, R. Pfannschmidt, F. Bruynooghe, B. Laugher, F. Bruhin 2004.

[18] R. Gautam, S. Vanga, F. Ariese, S. Umapathy, “Review of multidimensional data processing approaches for Raman and infrared spectroscopy” EPJ Tech. Instrum. 2015, 2, 1–38.

[19] J. R. Beattie, J. V. Glenn, M. E. Boulton, A. W. Stitt, J. J. McGarvey, “Effect of signal intensity normalization on the multivariate analysis of spectral data in complex ‘real-world’ datasets” J. Raman Spectrosc. 2009, 40, 429–435.

[20] X. Huang, J. Song, B. C. Yung, X. Huang, Y. Xiong, X. Chen, “Ratiometric optical nanoprobes enable accurate molecular detection and imaging” Chem. Soc. Rev. 2018, 47, 2873–2920.

[21] In Carbon Alloys, Elsevier Science, 2003, pp. 285–298.

[22] S. Zhang, Q. Liu, H. Zhang, R. Ma, K. Li, Y. Wu, B. J. Teppen, “Structural order evaluation and structural evolution of coal derived natural graphite during graphitization” Carbon 2020, 157, 714–723.

[23] S. K. Teh, W. Zheng, K. Y. Ho, M. Teh, K. G. Yeoh, Z. Huang, “Diagnostic potential of near-infrared Raman spectroscopy in the stomach: differentiating dysplasia from normal tissue” Br. J. Cancer 2008, 98, 457–465.

[24] P. Carmona, M. Molina, E. López-Tobar, A. Toledano, “Vibrational spectroscopic analysis of peripheral blood plasma of patients with Alzheimer’s disease” Anal. Bioanal. Chem. 2015, 407, 7747–7756.

[25] E. J. Haugen, A. K. Locke, L. H. Dao, A. B. Walter, P. K. Rasiah, J. S. Baba, M. A. Buendia, H. Correa, G. Hiremath, A. Mahadevan-Jansen, “Biochemical detection of pediatric eosinophilic esophagitis using high wavenumber Raman endoscopy and stimulated Raman microscopy” Sci. Rep. 2025, 15, 22471.

[26] P. Fu, Y. Zhang, S. Wang, X. Ye, Y. Wu, M. Yu, S. Zhu, H. J. Lee, D. Zhang, “INSPIRE: Single-beam probed complementary vibrational bioimaging” Sci. Adv. 2024, 10, eadm7687.

