## Supplementary Information for "msaGUI: Multispectral Analysis Graphical User Interface for Ratiometric Analysis and Background Correction"

**Statistics**
Below are mathematical representations of the statistics displayed below images. For each, *x* is the value of each pixel.

Count ($n$): *n* is the number of pixels in an image.

Mean ($\overline{x}$), where:

$$\overline{x} = \frac{\sum x}{n}$$

Median ($\tilde{x}$), where For an ordered set: $x_1 \leq x_2 \leq \cdots \leq x_n$:

$$\tilde{x} = \begin{cases} x_{\frac{n+1}{2}} & if\ odd \\ \frac{x_{\frac{n}{2}} + x_{\frac{n}{2}+1}}{2} & if\ even \end{cases}$$

Max ($x_{max}$):

$$x_{max} = \max(x_1, x_2, \ldots, x_n)$$

Standard Deviation (Std Dev, $\sigma$):

$$\sigma = \sqrt{\frac{1}{n}\sum_{i=1}^{n}(x_i - \bar{x})^2}$$

Standard Error (Std Error, $SE$):

$$\text{SE} = \frac{s}{\sqrt{n}}$$

```
10.97518,10.70196,10.8024,10.85626,10.56708,10.36459,10.30516
12.06819,14.11398,16.84036,19.26426,19.65212,20.58116,21.75483
12.93235,15.14095,17.89588,20.13307,20.6248,21.89983,22.77097
11.53448,14.8626,17.36334,19.77483,21.75422,22.34023,22.53391
11.09152,14.16566,16.90567,19.39143,21.96467,22.20977,22.51039
11.41531,14.33051,17.35577,19.48473,21.53819,23.51807,23.36234
11.3909,15.01118,17.96006,19.78769,21.30638,22.54708,22.44309
```

**Figure S1.** Example of how raw data is stored in a csv file.

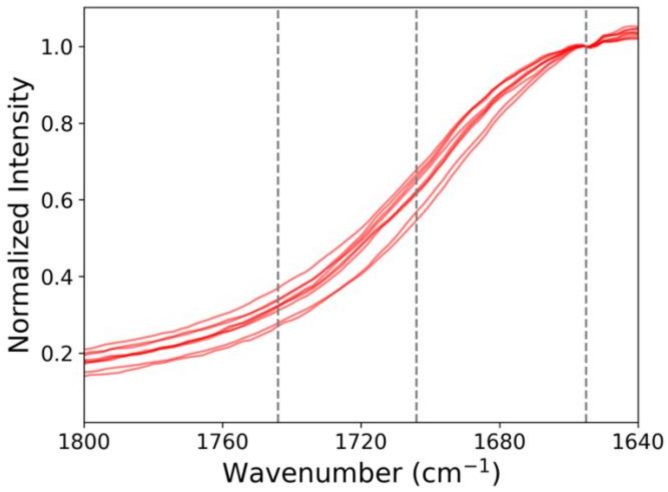


**Figure S2.** Spectra collected in the cytoplasm of Huh-7 cells fed with $^{12}$C glucose DMEM and normalized at 1655 cm$^{-1}$. Vertical lines drawn at 1655, 1703, and 1744 cm$^{-1}$ are used to determine the water and amide I (1655 cm$^{-1}$) background at the $^{13}$C=O (1703 cm$^{-1}$) and $^{12}$C=O (1744 cm$^{-1}$) ester carbonyl modes of cholesteryl ester and triglycerides. Using these correction factors, a water background image can be rescaled and subtracted from ester carbonyl images collected in lipid droplets.

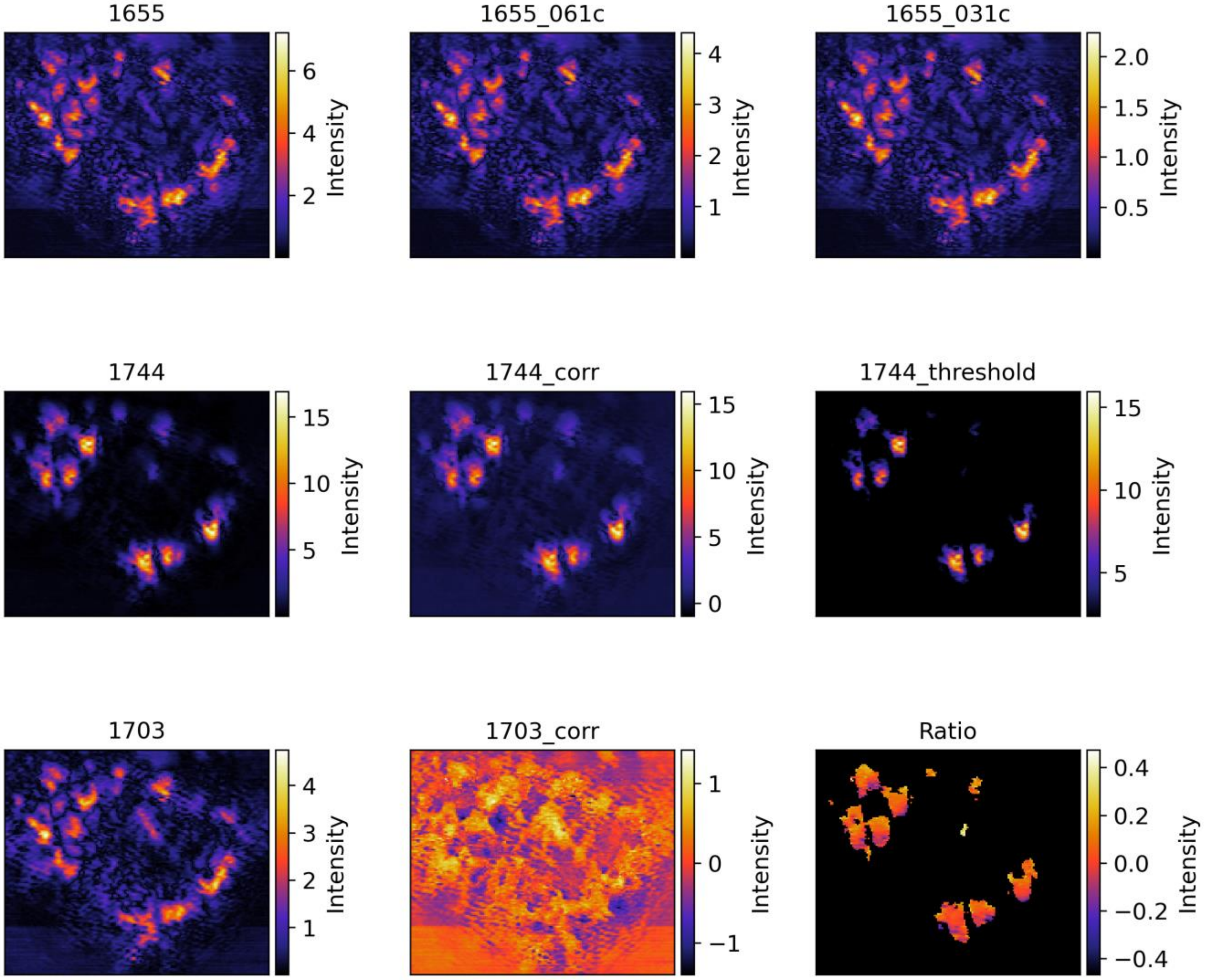


**Figure S3.** Group image generated from msaGUI.

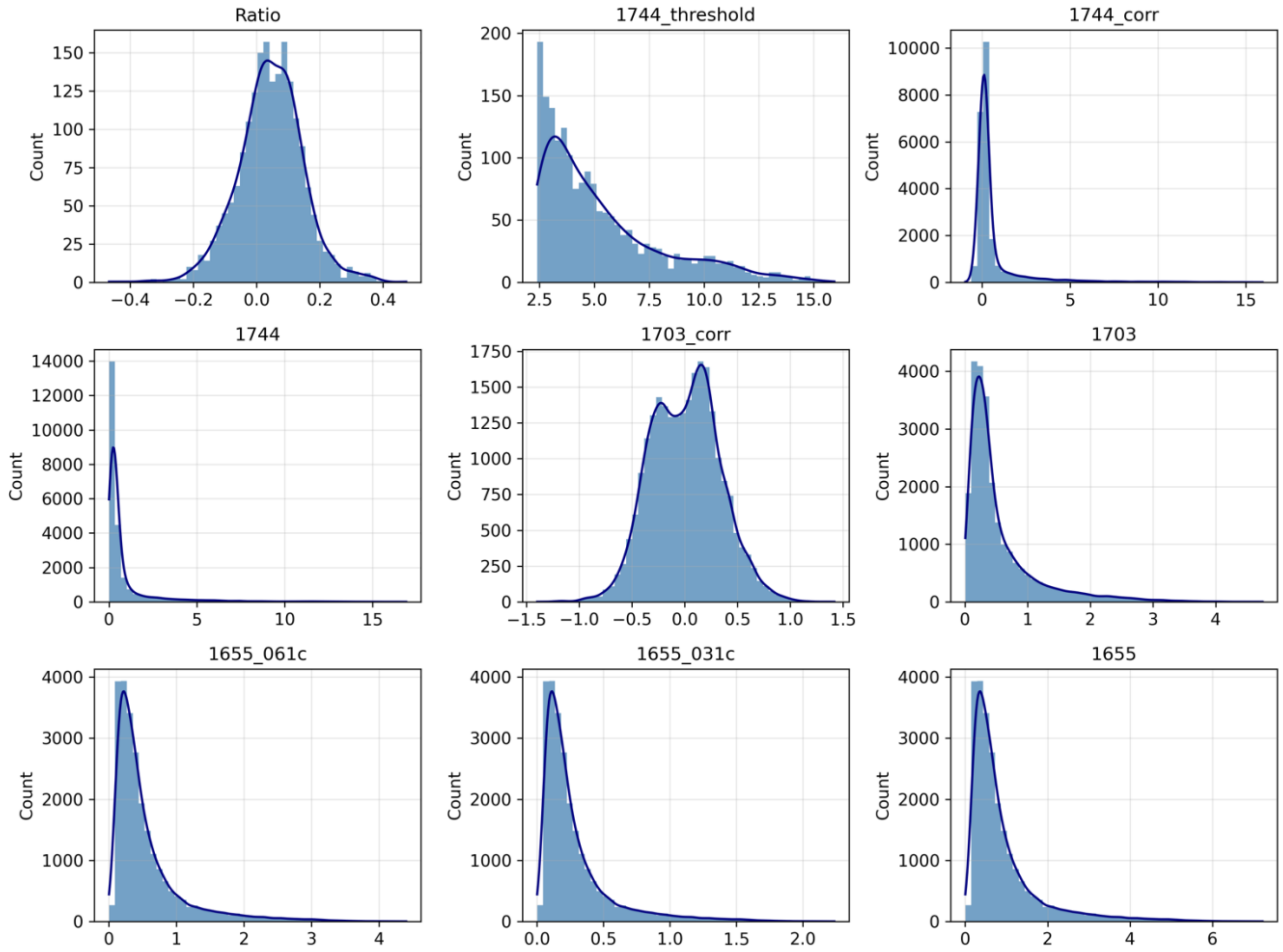


**Figure S4.** Group histogram generated from msaGUI.
